# A Multi-Agent Approach to Generating Context-Rich Gene Sets

**DOI:** 10.1101/2025.11.23.690073

**Authors:** Ebunoluwa Makinde, Farhad Maleki, Katie Ovens

## Abstract

Gene sets are collections of genes that share a common biological function, process, or component that can be used to get insight into the biological relevance of genomic data. Databases containing these gene sets aids in a wide array of analytical methods. The results of these methods, such as gene set analysis or phenotype-based gene prioritization, depend on the quality of the gene sets. Despite the extensive literature and genetic data available for constructing these databases, they often lack sufficient biological context. Current curation methods rely on labour-intensive expert manual curation from literature and datasets, as well as automated methods that are not context-aware. Therefore, there is a significant opportunity to utilize publicly available literature to bridge this gap and create more precise gene sets. With the advancement of natural language processing technologies, particularly large language models, this task can be performed more efficiently. In this work, we present a multi-agent system that utilizes the Llama 3, DeepSeek, and Qwen open-source large language models to analyze PubMed abstracts, allowing us to reconstruct gene sets in existing databases that better reflect specific biological contexts. Our approach consists of two pipelines. One verifies the inclusion of genes in a gene set by proof of evidence in the abstracts showing the association between the gene and the gene set. The second pipeline parses through the abstracts to identify genes not already included in the gene set for potential inclusion. To evaluate the proposed approach, we reconstructed a random selection of gene sets within the Human Ontology Phenotype (HPO). Our analysis shows that 149 of these gene sets have a similarity of 65.18% when compared to the original HPO gene sets, aligning well with the current HPO database. Additionally, we found an average of 3.15 new genes not included in the HPO gene sets, each supported by verified literature linking them to their respective gene sets. This highlights that our updated gene set database better reflects the current state of biological findings.

## 1 Introduction

Research into genes that contribute to various biomedical conditions has expanded rapidly, supported by advances in high-throughput omics technologies. These technologies, such as microarrays and RNA sequencing, are used to investigate the activity of thousands of genes in gene expression experiments, leading to an increase in available biological data [25,4]. The growing volume of biological data presents researchers with the challenge of extracting and interpreting biologically meaningful patterns. Gene lists generated from these experiments can provide insight; for example, genes expressed in cancer cells can indicate signaling pathways that suggest potential therapeutic approaches [14].

One way scientists have organized and interpreted these findings is through the grouping of genes with shared biological functions into gene sets. A gene set can be defined as a collection of genes that work together to perform a specific biological process, function, or component. These gene sets are compiled into gene set databases, which serve as structured repositories that enable comparative and interpretive biological analysis. Gene sets and related analytical methods have become essential tools for deriving meaning from genetic data, including approaches such as gene set analysis and phenotype-based gene prioritization. Gene set analysis is a powerful method used to extract meaningful insights into a range of biological conditions using high-throughput gene expression data. This method aims to assess whether experimental gene lists intersect with statistically and biologically relevant gene sets [25,2]. On the other hand, phenotype-based gene prioritization is used to identify and rank causative genes from a gene set database for observed phenotypes in a patient [15].

Both analytical approaches are heavily dependent on the availability and quality of well-curated gene set databases. Several widely used, literature-curated gene set databases exist, such as BioCarta [28], Gene Ontology (GO) [3], Human Phenotype Ontology (HPO) [9], Kyoto Encyclopedia of Genes and Genomes (KEGG) [16], and Reactome [6]. Each of these databases vary in scope, use case, and content. Gene sets and their corresponding databases play a crucial role in the outcomes of gene set analysis [26], and the reliability of phenotype-driven gene prioritization depends on the completeness and accuracy of these databases [15]. Although gene set databases are valuable resources, they often lack the specificity needed to capture the nuances of different biological contexts and genetic associations [27]. Curating more context-aware and biologically representative gene set databases can improve the identification of meaningful gene associations and understudied phenotypes [27]. With the extensive genomic data available in PubMed [30], there is an opportunity to apply data mining to create databases that better reflect biological complexity. The recent developments in artificial intelligence, particularly large language models (LLMs) and agentic systems, offers a means to automate and enhance this process.

The objective of this work is to reconstruct existing gene set databases and evaluate whether LLMs can effectively capture biological context to curate databases using only literature abstracts. Furthermore, this study will assess whether LLM-curated gene sets can surpass the quality of manually curated databases such as HPO. To achieve this, three open-source LLMs (Llama 3, Qwen, and DeepSeek) will be utilized to extract information from PubMed abstracts using Retrieval-Augmented Generation (RAG) techniques, with the goal of generating more accurate and biologically meaningful gene sets.

The remainder of this paper is organized as follows: Section 2 we review current methods for curating gene set databases and utilizing LLMs to improve them. We then analyze the novelty introduced by our research. Then, we explain our methodology for curating these context-rich gene sets using LLMs in a multiagent system in Section 3. In section 4, we describe the evaluation methods and the current results. Then we present our discussion of the quality of our LLM-curated gene set database in Section 5. Lastly, in Section 6, we conclude by presenting the summary of our findings and also pointing to future directions that can be taken to improve upon current gene set database curation and gene set analysis methods.

## 2 Background and Related Work

### 2.1 Large Language Models and Retrieval-Augmented Generation

Large language models (LLMs) are a subset of generative artificial intelligence where machine learning models are trained to identify patterns in data, enabling them to generate new data with similar properties to the training data [12]. These models have been adapted for various text processing tasks, such as information extraction, question answering, and summarization [14]. LLMs possess powerful mechanisms for understanding rich biological contexts [34]. Current research has produced several general-purpose LLMs, such as GPT-4 by OpenAI [1], Llama 3 by Meta [7], and Gemini by Google [33].

Despite being trained on extensive datasets, LLMs often lack context in specific domains. Studies have shown that performance can be improved for specific fields, such as medicine, through a process involving pretraining and fine-tuning [23]. However, these methods can be computationally expensive for ongoing use across various niche domains [29].

To address this issue, Retrieval-Augmented Generation (RAG) enhances a model’s knowledge base by retrieving information from external sources and incorporating it into prompts [20]. Augmenting the model in this way has been shown to substantially improve performance on domain-specific queries [23]. In our work, we will enhance the ability of three open-source LLMs to generate insights into the biological context of genes by incorporating external knowledge from PubMed abstracts using the RAG methodology.

### 2.2 Customary Approaches to Gene Set Database Curation

Gene Ontology (GO) is considered as a gold standard for gene set enrichment analysis tools [2]. GO annotations are generated through either manual or automated curation. Manual annotations are conducted by trained curators who extract information from full-text primary literature and functional data [13]. On the other hand, automated annotations employ algorithms that infer gene product functions based on orthology, domain composition, and sequence similarity [13]. Although automated approaches cover a broader annotation scope, manual curation provides higher precision and contextual relevance [13].

The Human Phenotype Ontology (HPO) provides a structured vocabulary of human phenotypic abnormalities and serves as a key resource for phenotype-driven gene prioritization and rare disease research [9,31]. Initial curation of HPO combined Java and Perl scripts to extract data from the OMIM database [31]. A process of both manual annotation and automated matching was then used to define HPO terms and annotate them with related disease concepts [31]. Over the years, the database has incorporated contributions from OrphaNet and DECIPHER to include more terms and annotations, this time including related genes [17,19,18]. Since its initial curation, the HPO has largely been collaborative, involving research groups from around the world that contribute to its terms and annotations [18].

An approach to improve gene set databases involves integrating multiple resources to create meta-databases such as the Molecular Signatures Database (MSigDB) [22]. MSigDB aggregates gene sets from diverse sources, including GO, HPO, Reactome, and KEGG, and contains one of the largest collections of gene sets available. Its curation process, similar to GO, involves both manual and computational means [22]. Automated methods identify overlaps between gene sets, while manual review ensures accurate annotation of biological themes, refinement of expression data, and precise classification of overlapping or redundant sets [21].

Manual curation of these databases introduces the possibility of human error, alongside tedious labour and redundant tasks, which can compromise the quality of gene sets. Furthermore, conventional computational methods used in these databases are limited in their ability to capture context-specific biological information. The present work seeks to address these limitations by reducing the manual workload involved in database curation and generating context-aware gene sets derived directly from published literature. By utilizing LLMs to automate and refine the curation process, this approach aims to enhance both the accuracy and biological relevance of gene set databases used in downstream analyses.

### 2.3 LLM-Based Approach for Generating Gene Sets and Insights for Gene Set Analysis

The tool *llm2geneset* proposed by Zhu et al., utilizes LLMs such as GPT-3.5, GPT-4o-mini, and GPT-4o to dynamically generate gene sets from natural language descriptions [38]. It uses a list of query genes, such as differentially expressed genes, optionally accompanied by biological context as input. The LLM is then queried to generate gene set descriptions based on this list of genes. These generated descriptions are then re-submitted to the model to produce corresponding gene sets by associating relevant genes with each description. The resulting gene sets are analyzed using Overrepresentation Analysis (ORA) to identify enriched sets. The method was benchmarked against human-curated databases, demonstrating substantial overlap [38]. Moreover, the performance of gene set analysis improved when using this pipeline compared with either ORA alone or direct LLM prompting.

In contrast, Hu et al. bypass the gene set curation stage and instead focus on how LLMs can be utilized for the functional interpretation of existing gene sets [12]. Their work introduces a pipeline in which an LLM is instructed to analyze a gene set, generate a biologically meaningful descriptive name, and evaluate its own performance on this task [12]. Results demonstrated that LLMs are capable of capturing the biological relevance and function of gene sets.

Building on Hu et al.’s work, Wang et al. propose Gene Agent—a language-based agent system designed to generate coherent biological process names for gene sets [34]. Gene Agent autonomously interacts with multiple domain-specific databases through Web APIs to gather contextual information from expert-curated resources [34]. Unlike previous methods, it incorporates a self-verification mechanism that evaluates and supports or refutes the LLM’s predictions. This design reduces hallucinations commonly observed in LLM outputs and refines generated annotations to improve the reliability of LLM-assisted gene set analysis.

#### Contrast with Proposed Technology

While innovative, the approach used by *llm2geneset* has significant room for improvement. One key limitation is its reliance on the pretrained knowledge of the LLM, which is neither comprehensive nor fully representative of current biomedical literature. In contrast, our proposed approach utilizes a Retrieval-Augmented Generation (RAG) methodology to ensure that curated gene sets are informed not only by pretrained knowledge of the model, but also by up-to-date information drawn directly from the current literature.

Hu et al.’s work provides a framework for evaluating how LLMs can generate biological insights from gene sets; however, it does not address the quality or specificity of the gene sets themselves. Our approach prioritizes this step, ensuring that gene sets are first curated to be biologically specific and contextually accurate before downstream inference is conducted, which was an aspect overlooked in Hu et al.’s study. Although Gene Agent implements a self-verification mechanism, it remains constrained by the limitations of its base model, GPT-4, which can produce biological process names that deviate from established references. By incorporating external domain-specific information through the RAG framework, our method enhances contextual grounding and mitigates hallucinations more effectively.

Furthermore, all three existing studies rely on proprietary OpenAI GPT models, which, despite their adaptability to diverse tasks, including gene-related analyses, have substantial costs and lack open-source accessibility. In contrast, our framework adopts three open-source models—Llama 3, Qwen 3, and DeepSeek R1—each demonstrating strong reasoning and generalization capabilities across multiple domains, making them suitable candidates for adaptation in this context [7,37,10].

## 3 Methods

The proposed framework consists of two systems designed for automated gene set curation. The first system functions as a gene set checker, serving as a verification pipeline that evaluates whether individual genes are appropriately included in a gene set based on supporting evidence extracted from the PubMed Abstract database. The second system operates as a gene set maker, utilizing the name of a gene set as an anchor to identify additional genes that may be biologically associated with it. Genes determined to be relevant are incorporated into the gene set—provided they are not already present—following validation through the gene set checker phase.

Each step involving a LLM was implemented using three selected open-source models: Llama 3.1 8B, DeepSeek R1 8B, and Qwen 3 32B. For this study, we reconstructed the Human Phenotype Ontology (HPO) database using this pipeline; however, the process is fully reproducible across other gene set databases such as Gene Ontology (GO) and Kyoto Encyclopedia of Genes and Genomes (KEGG). Figure 1 illustrates the overall workflow, outlining the sequential processes involved in extracting and organizing information from PubMed abstracts to curate gene set databases.

**Fig. 1.**
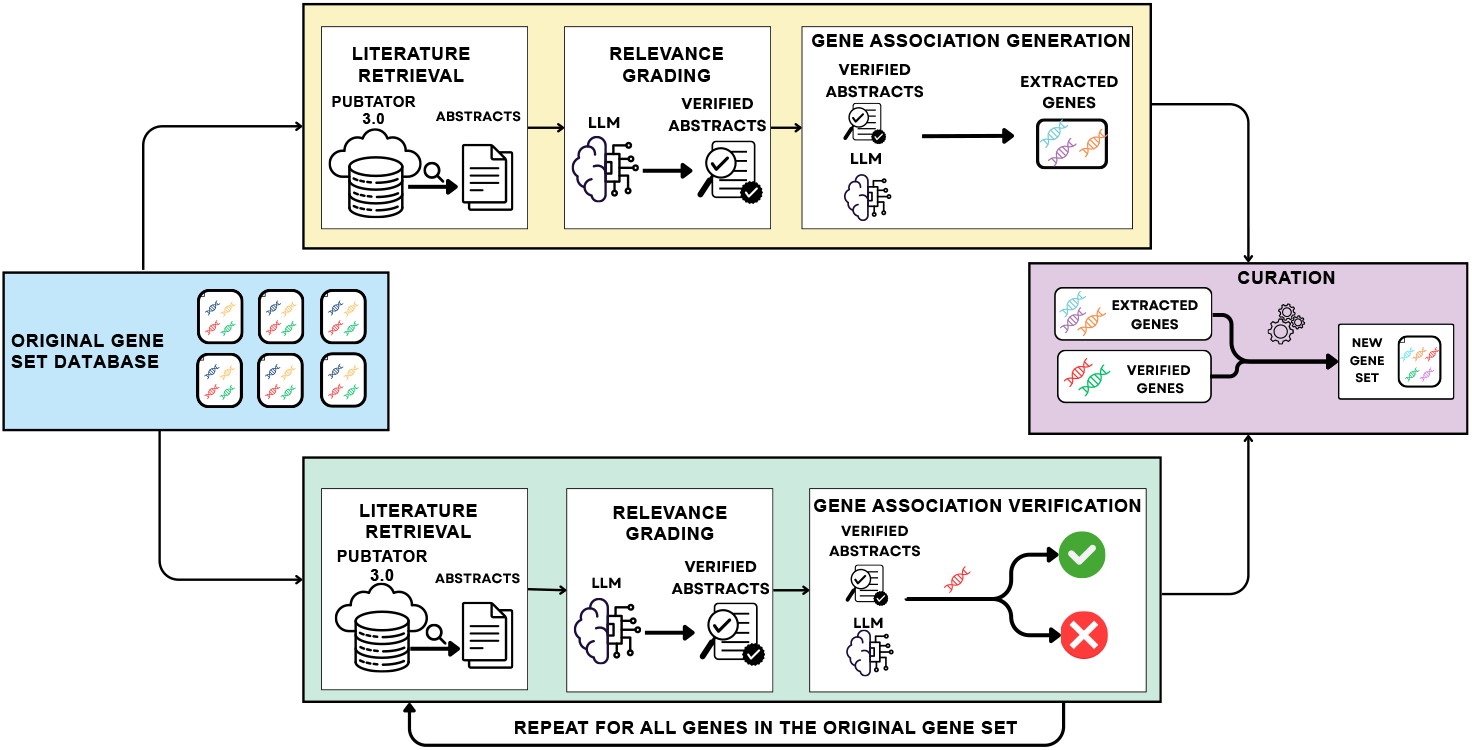
Overview of our gene set curation workflow. The framework takes gene set names from the original database and processes them through two pipelines. The Gene Set Maker pipeline (top) retrieves phenotype-related literature and extracts genes supported by verified abstracts, while the Gene Set Checker pipeline (bottom) validates gene–phenotype associations for genes already in the set. The curated outputs, consisting of newly extracted genes and verified existing genes, are combined to produce an updated literature-supported gene set.

### 3.1 Phenotype Extraction

Phenotype information was obtained by querying the HPO database API to extract the phenotype name, description, and corresponding synonyms. This is to get a broad context of what the phenotype entails to improve the semantic embedding when prompting the LLM, making it more phenotype-aware.

### 3.2 Literature Retrieval

For each extracted phenotype, the objective was to retrieve relevant PubMed abstracts that mention genes potentially associated with the phenotype. Rather than relying on PubMed’s default NCBI E-utilities API, we utilized PubTator 3.0, a biomedical text-mining system that supports advanced semantic and relation-based searches [35]. PubTator 3.0 provides annotations for PubMed abstracts covering entities such as genes, diseases, variants, chemicals, species, and cell lines. This enabled the retrieval of abstracts that were semantically relevant to the phenotype and contained explicit gene mentions.

When querying the system, we used the format “Phenotype Name” AND “genes.” This approach yielded a more concise list of abstracts that actually contained related genes, unlike simply searching by phenotype name, which often returned irrelevant results. Due to these streamlined queries, we limited our selection to the top 250 abstracts. This decision was driven by the need to manage storage and computational constraints, as collecting all returned abstracts would have led to a large number being discarded by the grader. Focusing on the top results ensured that we had a good subset containing the most associated genes related to the phenotype in question. After retrieving the top 250 abstracts or fewer, we then checked PubTator’s annotations to verify the presence of genes, filtering out any abstracts that did not contain genes.

For the gene set checker pipeline, for each gene associated with a phenotype, we send a query to PubTator 3.0 with the phenotype and the gene name in gene symbols format. Instead of submitting a regular search query, we use PubTator’s prefix for identifying genes, which is “@GENE_”, and append this to the gene before querying the pipeline with the modified gene and phenotype name. We only extract the first 10 abstracts. In a study of related entities, PubTator 3.0, through manual verification, showed a precision of 90% for the top 20 articles retrieved [35]. Therefore, selecting the top 10 would likely demonstrate an even greater correlation, and this approach also conserves computational time.

### 3.3 Relevance Grading

Before proceeding to the gene extraction step, it was necessary to address the issue that not all retrieved abstracts mentioning genes were truly indicative of a gene–phenotype association. Some abstracts referenced genes in contexts unrelated to the phenotype or described negative correlations, such as instances where individuals exhibiting the phenotype lacked expression of the gene. To ensure contextual accuracy and minimize hallucinations in subsequent stages, we first used the LLM as a grader.

The LLM was provided with the phenotype’s name, description, and synonyms and instructed to evaluate the relevance of each abstract by determining whether it described genes genuinely associated with the phenotype. The complete prompt used for this grading task is included in the Appendix. The model produced a JSON output containing a single key, binary_score, with one of two possible values (“yes” or “no”) indicating whether the abstract was deemed relevant.

The same grading process was applied in the gene set checker pipeline, with the evaluation criterion expanded to verify whether abstracts explicitly discussed associations between the phenotype and the gene. This additional filtering step ensured greater specificity and reliability in identifying valid gene–phenotype relationships for downstream curation.

### 3.4 Gene Association Generation

Once the filtered set of abstracts that mention genes to describe correlations with the phenotype is selected, the LLM is prompted again. In this phase, the model reviews each abstract alongside the phenotype’s metadata, which includes its name, synonyms, and description. It then extracts genes associated with the phenotype. The model returns a list of JSON objects, where each object includes the gene name, the supporting text excerpt (directly quoted from the abstract), the PubMed ID (PMID), and, when available, the journal source. The following JSON schema defines the structure of the gene set maker output:

~~~
[
  {
   “Gene”: “string”,
   “Source Reference”: “string”,
   “PMID”: “string | integer”,
   “Journal”: “string”
  }
]
~~~

For the gene set checker, the LLM produces a single JSON object confirming the presence or absence of the specified gene within the phenotype context. The corresponding JSON schema for this output is shown below:

~~~
{
  “Gene”: “string”,
  “Validation”: “string (yes | no)”,
  “Supporting Extract”: “string”,
  “PMIDS”: [“integer”]
}
~~~

### 3.5 Curation

In the curation phase, we consolidate the results produced by each LLM by merging newly identified genes and verified genes into a standardized Gene Matrix Transposed (GMT) file. To ensure identifier consistency, we use the MyGene Python library [36] to convert all gene symbols into their corresponding Entrez Gene IDs. Since gene symbols typically consist of short alphanumeric abbreviations, Entrez IDs provide numeric identifiers that are unique and unambiguous [24]. This conversion also serves as an additional validation step, as it enables cross-checking to ensure that the LLM has not incorrectly identified proteins or other biological entities as genes. The curated information is stored in an interactive HTML table generated with JavaScript, containing the phenotype name, source type (extracted, verified, or both), gene name, supporting abstract excerpt, and linked PubMed ID. This design allows for manual verification, ensuring transparency and reproducibility in gene inclusion decisions.

Each LLM generates its own curated gene sets, which are merged to produce a unified result. To increase confidence in gene inclusion, we apply a majority-voting strategy: a gene is included in the final set if it appears in at least two of the three LLM-generated outputs. The resulting gene sets are reported as the final curated output of the pipeline.

### 3.6 Comparison Metrics

To determine agreeability between our curated gene sets and what is currently curated, we use the Jaccard index to quantify similarity between two individual gene sets *G*_*i*_ and *G*_*j*_ as defined below [26]:

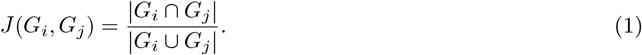

To quantify the similarity between entire gene set databases, we considered the union of all genes appearing in each database. Let 𝔾_1_ and 𝔾_2_ denote two gene set databases with gene sets

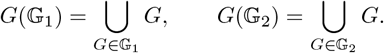

The overall database similarity is then defined as the Jaccard index over these aggregated gene sets:

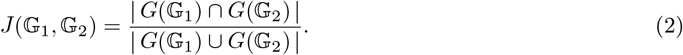

The Jaccard index ranges between 0 and 1, where 1 indicates identical gene content; in this work, we report it as a percentage for interpretability. Our source code as well as final outputs and datasets can be found here: https://github.com/EbunMak/multi-agent-generator-gene-set

## 4 Results

### 4.1 Evaluation of Curated Gene Set Metrics

In order to validate the continuity of our pipeline, we first curated a selection of 149 gene sets; the “_New” suffix will be used to indicate reconstructed databases by our pipeline. We then compared their agreement with their counterparts in the original HPO gene set database. To achieve this, we used Equation 2 to quantify similarity between two individual gene sets. This gave us a 65.18% similarity score when expressed as a percentage; this is what we define as overall database similarity. To help us understand how many of the total genes are shared between individual gene sets in both databases, we also calculated the mean similarity of each gene set (defined in Equation 1, resulting in 46.78%. In this subset, we also noted an average of 3.14 new genes added per gene set.

Evaluating solely this subset, we see that based on the similarity, there is agreement between HPO and HPO_New. With these initial results, we furthered our pipeline to reconstruct a total of 437 gene sets, including the 149 we first created. Table 1 summarizes an array of metrics obtained for these 437 gene sets when compared to the same 437 found in the original HPO database. As it reports, 3.41 new genes were added on average. This closely aligns with what we found for the smaller subset we initially curated. However, for this larger set, we also discovered an overall database similarity of 83.59%, which suggests that as more gene sets were curated, there was closer similarity between ours and the original.

**Table 1.** Summary of Gene Set Database Comparison Metrics.

| Metric | Value |
| --- | --- |
| Mean new genes per gene set | 3.41 |
| Mean lost genes per gene set | 54.42 |
| Mean % of lost genes | 34.58% |
| Maximum lost genes in a gene set | 1226 |
| Maximum new genes in a gene set | 31 |
| Maximum % of lost genes | 97.02% |
| Overall database similarity | 83.59% |
| Mean Jaccard similarity across gene sets | 57.66% |
| Total unique genes in HPO_New | 4823 |
| Total unique genes in HPO | 4909 |
| Shared genes across databases | 4476 |
| Total new genes in HPO_New not in HPO | 433 |
| Total genes in HPO not found in HPO_New | 347 |

#### Examining the Similarity Distribution among Gene Sets

The mean Jaccard similarity across gene sets was 57.66%. Figure 2a displays the distribution of the gene set similarity scores, all evaluated according to Equation 1 and expressed as percentages. The majority of the phenotypes exhibited a moderate to high overlap of 40% to 85%. The most common range was between 60% and 70%, indicating that a significant number of genes are shared across the gene sets of HPO and HPO_New. Overall, this shows that the curated HPO_New demonstrates agreement with HPO.

**Fig. 2.**
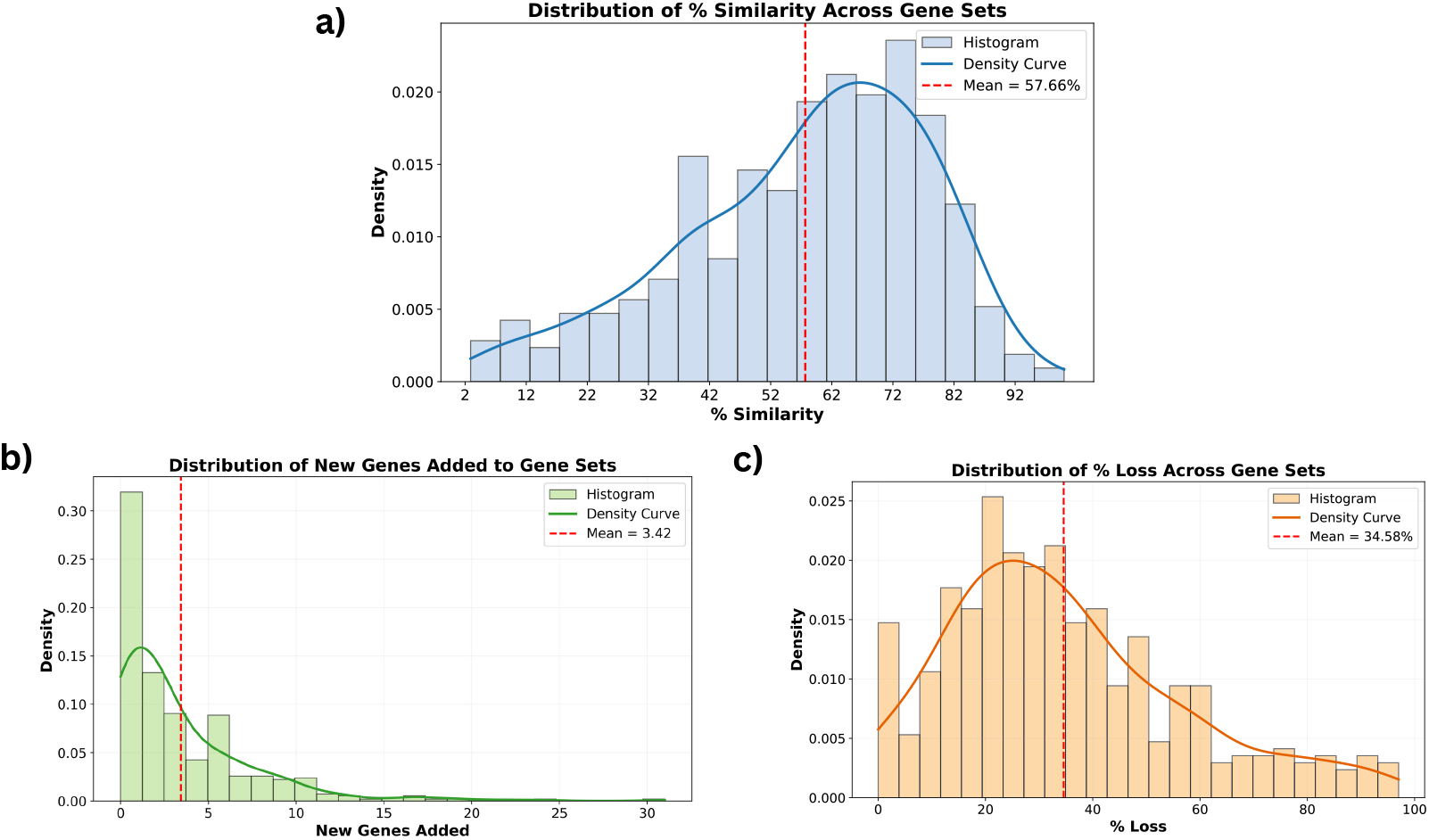
Distributions of gene set similarity and changes between databases. a) Percent similarity across gene sets. b) Number of newly added genes per gene set. c) Percent gene loss relative to original set size.

#### Examining the Distribution of New Genes Added

An average of 3.41 genes were added per gene set, with up to 31 new genes added for the phenotype, Abnormal metabolism. Figure 2b illustrates the distribution of new genes, where most gene sets had fewer than 2 genes added to them. Most genes remained stable; there were 21.74% of the genes (95 genes) that remained consistent with no new genes added, while the remaining 342 gene sets, for which new genes were added, suggest that some new biological insights were discovered merely from the literature. There were also two gene sets: Aplasia of the ulna and Patchy changes of bone mineral density that were completely the same between HPO_New and HPO.

#### Examining the Distribution of Lost Genes

Gene sets across a database vary widely in their original size, therefore we report percentage loss rather than absolute loss. This allows for changes to be interpreted relative to the scale of each gene set and enabling more meaningful comparison across HPO terms. Figure 2c, shows the distribution of the percentage loss is right-skewed, with the mean percent loss at 34.58% (dashed red line), indicating that on average, our pipeline includes 65.42% of HPO-annotated genes for each term. The majority of the gene sets exhibit moderate loss as 110 (25.17%) gene sets have low loss less than 20%, 175 (40.05%) with loss of 20-40%, and 65 (14.18%) with high loss (greater than 60%) These findings indicate that the curation pipeline verifies and retrieves a good amount of gene annotations documented in HPO. A minority of terms are marked by high percent loss, as reflected in the long-tail of the distribution. We also note that there are no gene sets for which all original genes were completely lost.

### 4.2 Case Study: Abnormal Metabolism

To evaluate whether the newly added genes generated by our pipeline were truly contextually relevant, we manually validated each gene added across the top ten phenotypes. The complete validation table is provided in the Appendix. Here, we focus on the phenotype Abnormal Metabolism, which had the highest number of newly added genes (31).

#### Genes Extracted From PMID: 34616452 [5]

The genes *MDK, SLC1A1, SGCB, MALAT1, PILRB, IGHG1, FZD1, IFITM1, MUC20, KRT80*, and *SALL1* were extracted by our pipeline as novel genes associated with Abnormal Metabolism. Both Llama 3.1 and DeepSeek R-1 independently identified these genes as metabolism-related based solely on the abstract. Importantly, all eleven genes trace back to the same source abstract (PMID: 34616452). A sample of the DeepSeek evidence is:

~~~
{
  “Gene”: “MDK”,
  “Source Reference”: “The Riskscore model was obtained by multiplying the expression levels of these 12 genes with the corresponding coefficients of the multivariate regression.”,
  “PMID”: “34616452”,
  “Journal”: “J Oncol”
}
~~~

A sample of Llama 3.1’s response is included below, but one thing to note is that it assigned an incorrect PMID which is something we also observe in other responses:

~~~
{
  “Gene”: “SLC1A1”,
  “Source Reference”: “Identification of Prognostic Metabolism-Related Genes in Clear Cell Renal Cell Carcinoma.”,
  “PMID”: “37563740”,
  “Journal”: “J Oncol”
}
~~~

Despite the incorrect PMID, the supporting reference still corresponds to the same abstract. The abstract explicitly states: *“Finally, 12 differentially expressed genes (MDK, SLC1A1, SGCB, C4orf3, MALAT1, PILRB, IGHG1, FZD1, IFITM1, MUC20, KRT80, and SALL1) were filtered out using the random forest model*.*”* The study focuses on clear cell renal cell carcinoma (ccRCC), a cancer described as having abnormal metabolism, and notes: *“The Riskscore model was associated with most of the metabolic processes*.*”* Because these genes are directly tied to metabolic dysregulation in ccRCC, their addition to the Abnormal Metabolism gene set is therefore justified.

#### Genes Extracted From PMID: 38025761 [32]

The 14 genes *DLAT, SEPHS1, ACADS, UCK2, GOT2, ADH4, LDHA, ME1, TXNRD1, B4GALT2, AK2, PTDSS2, CSAD*, and *AMD1* were reported by Llama 3.1 and Qwen 3 as associated with Abnormal Metabolism, while DeepSeek R-1 identified only *GOT2* from the same abstract. All three models ultimately pointed to the same source abstract (PMID: 38025761), with representative extractions shown below. Llama 3.1:

~~~
{
  “Gene”: “GOT2”,
  “Source Reference”: “A novel metabolism-related gene signature in patients with hepatocellular carcinoma.”,
  “PMID”: “38025761”,
  “Journal”: “PeerJ”
}
~~~

Qwen 3:

~~~
{
  “Gene”: “AMD1”,
  “Source Reference”: “A Metabolism-Related Risk Score (MRRS) model was constructed using 14 MRGs (DLAT, SEPHS1, ACADS, UCK2, GOT2, ADH4, LDHA, ME1, TXNRD1, B4GALT2, AK2, PTDSS2, CSAD, and AMD1).”,
  “PMID”: “38025761”,
  “Journal”: “PeerJ”
}
~~~

The abstract mentions that hepatocellular carcinoma (HCC) is associated with abnormal metabolism and that these genes are explicitly used to construct a metabolic risk score. Thus, all 14 genes are valid additions to the Abnormal Metabolism phenotype.

#### Genes Extracted From PMID: 36809279 [11]

Qwen and Llama 3.1 extracted the genes *SIRT3, SIRT4, SIRT5, SOD1, SOD2*, and PARP1 from the abstract describing the oncometabolic role of mitochondrial sirtuins in glioma patients. Qwen provided the detailed supporting evidence: *“Results analysis showed significant down-regulation of SIRT4 (p = 0*.*0337), SIRT5 (p < 0*.*0001), GDH (p = 0*.*0305), OGG1-2alpha (p = 0*.*0001), SOD1 (p < 0*.*0001) and SOD2 (p < 0*.*0001) in glioma patients*.*”* The study demonstrates that mitochondrial metabolic enzymes and oxidative stress regulators (*SIRT3, SIRT4, SIRT5, SOD1, SOD2*) and DNA repair–linked *PARP1* show dysregulation in glioma, all of which are consistent with the metabolic phenotype.

#### Genes Extracted From PMID: 31733337 [8]

Qwen and Llama 3.1 extracted these two genes (*NFE2L1, NGLY1*) from a study on hepatocyte-specific NGLY1 deficiency. Qwen again provided the correct PMID and accurate supporting text: *“hepatocyte-specific Ngly1-deficient mice showed abnormal hepatocyte nuclear size/morphology with aging*…*”* and *“We showed that the processing and localization of the transcription factor, nuclear factor erythroid 2-like 1 (Nfe2l1), was impaired in the Ngly1-deficient hepatocytes*.*”* NGLY1 deficiency disrupts NFE2L1 processing, leading to abnormal lipid and metabolic responses in hepatocytes. Because this metabolic impairment is directly evidenced in the abstract, both genes are valid additions.

Overall, we observed that manual validation confirmed that 31 newly extracted genes had clear evidence of inclusion in abstracts linking them to Abnormal Metabolism. We also observed that Qwen 3 provided the most reliable PMIDs and clearest direct evidence. The model Llama 3.1 captured relationships but often mismatched PMIDs or provided incomplete citations. DeepSeek R-1 reported fewer genes but was valuable in confirming gene inclusion.

## 5 Discussion

Literature is being updated every day, and the data available is becoming more and more accessible. However, traditional approaches to curating gene set databases always lag behind and fall short of keeping up with the current literature, or rely on automated methods not able to decipher literature well enough to get the full context. Our approach is able to index existing literature, updating these databases to better capture the reality of biomedical research. We demonstrate this by showing that, on average, 3.41 genes might be missing from the gene sets in the HPO database, and our pipeline can extract them. Specifically, we show that the gene set for which the most new genes were added, abnormal metabolism, had all the genes verified by the abstracts the models return for verification.

Our model, however, is not completely agnostic to the current state of the art, with a similarity in the genes contained of 83.59% and the loss of original gene sets being an average of 34.58%, supporting the capacity of the pipeline for effective contextual inference and verifying that what is in the HPO database correctly aligns with the literature. However, we acknowledge that there is a subset within our curated gene sets that demonstrates high loss. This could stem from the fact that gene set databases contain many genes that should not be there, as they do not capture the biological context of the gene sets, have variation, and redundancy [21]. By having two or more LLMs verify the presence of genes based on existing scientific proof creates a superior method that filters out genes that ought not to be included in the gene sets, hence capturing the true essence of the biological associations of gene sets. We also account for the possibility that the genes not mentioned could also be a result of not parsing through enough abstracts in the literature, or that these gene annotations reported in HPO are not found in what is available on PubMed, as it is only one database and not including full-text articles.

Notably, gene set databases are only as useful as their ability to provide accurate answers through gene set analysis or phenotype-based gene prioritization. With our model being able to infer new genes, both methods can be substantially improved. For instance, the new genes enhance the precision of gene set analysis, allowing for the more accurate identification of differential gene sets. For phenotype-based gene prioritization, in a clinical context, this directly enhances the ability to prioritize disease-relevant genes better. Such improvements are particularly valuable for rare disease diagnosis, where capturing recently discovered or less studied gene-phenotype relationships can be critical for interpreting patient data and finding accurate diagnoses.

That being said, our workflow is time and computationally expensive. Future work should consider parallelizing pipelines and optimizing prompts to reduce the number of generations required. This would also allow for multiple reruns to minimize variability and ensure the model provides consistent results. We also experience a myriad of challenges with the LLM’s generations, from broken returned JSONs to false PMID associations. Further work should involve experimenting with the optimal number of documents necessary for generating high-quality gene sets. There should also be more checks and methods implemented, such as better prompt engineering, to ensure the LLM generations are consistent. Lastly, this work should extend to recreate the full collection of gene sets on HPO and even more commonly used for gene set analysis, like GO. Consequently, the reconstructed larger, more extensively used gene sets should be employed for conducting gene set analysis and phenotype-based gene prioritization, and their performance should be compared against the original versions. This will serve as a better test of the pipeline’s quality, establishing it as a superior curation method.

## 6 Conclusion

In conclusion, the choice of curated gene set databases plays a significant role in determining the outcomes of gene set analysis and phenotype-based gene prioritization. Therefore, our objective was to provide a gene set database rich in biological context to enhance these analysis outcomes. While efforts have been made to curate novel gene set databases, these do not provide comprehensive solutions for curating insightful gene sets based on the current literature. Thus, we present a multi-agent system setup that integrates external genetic knowledge into LLMs via RAG, elevating the quality of gene set databases. We have demonstrated that this pipeline is capable of curating gene sets that are not only similar to the original gene sets but also capture more information that is absent in the original version. It also saves time and minimizes the labour associated with manual curation and overcomes the shortcomings of the primitive computational methods used to curate the original database version. By utilizing LLMs and the vast sea of scientific literature, we present a new method capable of enhancing the quality of current gene sets. Its integration holds the potential to profoundly unlock new genetic insights.

