## Appendix for "A Multi-Agent Approach to Generating Context-Rich Gene Sets"

### Appendix: Supplementary Materials

#### Pipeline Implementation Details

**A. Phenotype Extraction** We downloaded the `phenotype_to_gene.txt` file from the Human Phenotype Ontology (HPO) database, which lists one phenotype–gene association per line. This file is the source from which MSigDB's HPO gene set database is curated. While MSigDB contains 5,748 curated HPO gene sets, the original file contains more associations. We reconstructed the data so that each line corresponds to a phenotype together with the full set of its associated genes.

Next, we identified the intersection between phenotypes present in MSigDB and those listed in the HPO file, yielding 5,567 phenotypes for reconstruction. Using our phenotype extractor pipeline, we queried the HPO API to retrieve phenotype identifiers, definitions, and synonyms. All phenotype metadata were stored as a list of JSON objects with a fixed schema.

**B. Literature Retrieval** Each phenotype JSON entry was processed through the retrieval pipeline. For every phenotype, we queried the PubTator 3.0 API using:

```
{phenotype name} AND genes
```

The query returns a list of PubMed IDs (PMIDs). We selected the first 250 PMIDs and then queried a second PubTator endpoint that exports full annotations for each publication, including abstracts and gene mentions. Only abstracts containing at least one annotated gene were retained. Retrieved abstracts were saved in phenotype-specific JSON files to enable persistence and avoid redundant re-querying.

**C. Abstract Relevance Grading** Abstract relevance grading was performed independently for each of the evaluated models (DeepSeek-R1 8B, Qwen3-32B, and Llama-3.1-8B) using the following instruction:

```
You are a scientific grader assessing the relevance of retrieved scientific
abstracts to a biological or disease-related question.

If the abstract contains keywords or semantic meaning related to the question,
grade it as relevant. Otherwise, mark it as not relevant.

Respond ONLY with:
{"binary_score": "yes"} or {"binary_score": "no"}.
```

Filtered abstracts (graded as "yes") were stored separately for each model.

**D. Gene Extraction** Gene extraction was run separately for each model. For each phenotype, filtered abstracts were reformatted to include the PMID, journal (if available), title, and abstract text. The model was then prompted with:

```
You are an assistant for a gene set curation task.

Your role is to identify genes possibly related to the biological process or
disease described in:
{question}

using the following context:
{context}

Respond ONLY with a list of JSON objects with keys:
1. "Gene"
2. "Source Reference"
```

```
3. "PMID"
4. "Journal"

Ensure that:
- No genes are duplicated.
- Each "Source Reference" comes from the corresponding PMID.
- Output is valid JSON only.
```

The question string used was:

```
Identify genes associated with phenotype "<name>",
definition: "<definition>".
```

This step completes the maker pipeline, and all extracted genes were stored in JSON format.

**E. Gene Checker Pipeline** After completing the maker pipeline for a phenotype, we executed the checker pipeline. For each gene in the original HPO gene set, PubTator 3.0 was queried for the top ten abstracts mentioning both the phenotype and the gene. Retrieved abstracts were stored in phenotype- and gene-specific JSON files.

**E.1. Abstract Relevance Grading** The following instruction was used:

```
You are a biomedical research assistant.

Determine whether an abstract discusses BOTH:
1. The specified phenotype, and
2. The specified gene.

Return ONLY:
{"binary_score": "yes" or "no"}.
```

**E.2. Gene Association Verification** We validated gene–phenotype associations using the prompt:

```
You are an assistant for a gene set validation task.

Context:
{context}

Question:
{question}

Return JSON with:
1. Gene
2. Validation ("yes" or "no")
3. Supporting Extract (direct quotes; 5 lines if multiple abstracts)
4. PMIDS (at least one PMID if validation="yes")
```

Both steps were conducted independently for each model.

**F. Consolidation and Gene Set Reconstruction** All extracted and validated genes were aggregated through a final curation pipeline. For each model, we generated:

- a GMT file containing the consolidated gene set, and
- an accompanying HTML table for inspection.

Sample output files are available in the `out/genesets` directory of our GitHub Repository.

A final consensus GMT file was constructed by including a gene in a phenotype's gene set if at least two of the three models identified or validated the gene.

#### Manual Verification of Gene Sets

| Gene(s) | Supporting PMID | LLM(s) | Validation |
|---|---|---|---|
| MDK, SLC1A1, SGCB, MALAT1, PILRB, IGHG1, FZD1, IFITM1, MUC20, KRT80, SALL1 | 34616452 | Llama 3.1, DeepSeek R-1 | valid |
| DLAT, SEPHS1, ACADS, UCK2, GOT2, ADH4, LDHA, ME1, TXNRD1, B4GALT2, AK2, PTDSS2, CSAD, AMD1 | 38025761 | Llama 3.1, Qwen 3 | valid |
| SIRT3, SIRT4, SIRT5, SOD1, SOD2, PARP1 | 36809279 | Llama 3.1, Qwen 3 | valid |
| NFE2L1, NGLY1 | 31733337 | Llama 3.1, Qwen 3 | valid |

**Table 2.** Manual Validation for Phenotype: Abnormal Metabolism

| Gene(s) | Supporting PMID | LLM(s) | Validation |
|---|---|---|---|
| ACTN4, CAPN12, ECH1, EIF3K, LGALS4, LGALS7, LGALS7B, MAP4K1, SYNPO2 | 37131961 | Qwen 3, DeepSeek R-1 | valid |
| AGPAT2 | 36978948 | Llama 3.1, DeepSeek R-1 | invald |
| BSCL2 | 34409079, 38525541 | Llama 3.1, Qwen 3, DeepSeek R-1 | invalid |
| CD19 | 35169706 | Llama 3.1, Qwen 3, DeepSeek R-1 | invalid |
| CD81 | 34359359, 36605192, 37251406, 39873880, 40297314, 40661769, 40707916 | Llama 3.1, DeepSeek R-1 | invalid |
| CR2, IDS, MS4A1 | | Llama 3.1, DeepSeek R-1 | invalid |
| ICOS | 37705722 | Llama 3.1, Qwen 3, DeepSeek R-1 | invalid |
| IRF2BP2 | 40531845 | Qwen 3, DeepSeek R-1 | invald |
| NFKB1 | 32234376, 34572989, 35497360 | Llama 3.1, Qwen 3, DeepSeek R-1 | invald |
| NFKB2 | 32184784, 37743226, 38366068 | Llama 3.1, Qwen 3, DeepSeek R-1 | invald |
| TNFRSF13B | 32991402, 39981695 | Llama 3.1, DeepSeek R-1 | invald |
| TNFSF12 | 34172716, 35498539, 39905000 | Llama 3.1, Qwen 3, DeepSeek R-1 | invalid |

**Table 3.** Manual Validation for Phenotype: Abnormal vascular morphology

| Gene(s) | Supporting PMID | LLM(s) | Validation |
|---|---|---|---|
| ANXA5, CD44, CDCA8, COL6A1, NCAM1, PLK4, SPP1 | 35884419 | Llama 3.1, Qwen 3, DeepSeek R-1 | invalid |
| CASP8 | 35928267 | Qwen 3, DeepSeek R-1 | invalid |
| CCL2 | 33401355, 34000383, 35461349 | Llama 3.1, Qwen 3, DeepSeek R-1 | valid |
| CDT1, HJURP, TOP2A, VEGFA | 35884419 | Qwen 3, DeepSeek R-1 | invalid |
| CLDN4 | 35742959 | Llama 3.1, Qwen 3, DeepSeek R-1 | invalid |
| CX3CL1 | 34671562 | Qwen 3, DeepSeek R-1 | invalid |
| EPHB4 | | Llama 3.1, DeepSeek R-1 | valid |
| FOXM1 | 35884419 | Llama 3.1, Qwen 3 | invalid |
| RARB | 34203310, 34238300, 37627010 | Llama 3.1, Qwen 3, DeepSeek R-1 | valid |
| SLC2A1 | 38169393 | Llama 3.1, DeepSeek R-1 | invalid |

**Table 4.** Manual Validation for Phenotype: Abnormality of the bladder

| Gene(s) | Supporting PMID | LLM(s) | Validation |
|---|---|---|---|
| ACTA2, ACTG2, DKK1, DKKL1, GPC3, GREM1, KCNMB1, KCNMB2, MYL9, PPP1R12B, RSPO3, SFRP5, TAGLN, WNT10B | 38473810 | Llama 3.1, Qwen 3 | valid |
| BAX, BCL2 | 32653936 | Llama 3.1, Qwen 3 | invalid |
| CCL2, CXCL12, FOS | 32684870 | Llama 3.1, Qwen 3 | valid |
| MCL1 | 36510562 | Llama 3.1, Qwen 3 | valid |

**Table 5.** Manual Validation for Phenotype: Abnormality of the intrinsic pathway

| Gene(s) | Supporting PMID | LLM(s) | Validation |
|---|---|---|---|
| CD6 | 36275816 | Llama 3.1, Qwen 3, DeepSeek R-1 | valid |
| COA4, GAPDH, GSTP1, HAGH, MFF, PHB2, TSPOAP1 | 37082114 | Llama 3.1, Qwen 3, DeepSeek R-1 | valid |
| CRYAB | 37196088, 39444000, 40372913 | Llama 3.1, Qwen 3, DeepSeek R-1 | valid |
| FMR1 | 34789272 | Llama 3.1, Qwen 3, DeepSeek R-1 | valid |
| GFAP | 34799461 | Llama 3.1, Qwen 3 | valid |
| GSDMB, IKZF3, ORMDL3, PHF20L1, ZPBP2 | 37478401 | Llama 3.1, DeepSeek R-1 | invalid |
| IRF2BP2 | 35795667, 37350971, 39059757 | Llama 3.1, Qwen 3, DeepSeek R-1 | valid |
| ITPR3 | 39560673 | Llama 3.1, DeepSeek R-1 | valid |
| MTHFR, PADI4, TRAF1 | 37797401 | Llama 3.1, DeepSeek R-1 | invalid |
| RNASET2 | 38960478 | Llama 3.1, DeepSeek R-1 | valid |
| TMEM230 | 38960478 | Llama 3.1, Qwen 3, DeepSeek R-1 | valid |
| TNFSF12 | 34542797, 34716660 | Llama 3.1, Qwen 3, DeepSeek R-1 | valid |

**Table 6.** Manual Validation for Phenotype: Autoimmunity

| Gene(s) | Supporting PMID | LLM(s) | Validation |
|---|---|---|---|
| ADGRV1, GRHL2, GRXCR2, MYO6, OTOF, OTOG, PCDH15, PDZD7, POU3F4 | 32048449 | Llama 3.1, Qwen 3 | valid |
| ANKRD11 | 40760574 | Qwen 3, DeepSeek R-1 | valid |
| ATP2B2 | 30535804 | Llama 3.1, Qwen 3, DeepSeek R-1 | valid |
| EYA4 | 35578334 | Llama 3.1, Qwen 3, DeepSeek R-1 | valid |
| FDXR | 39780253 | Qwen 3, DeepSeek R-1 | valid |
| SMPX | 33708524 | Llama 3.1, Qwen 3, DeepSeek R-1 | valid |
| TMPRSS3 | 37713394 | Qwen 3, DeepSeek R-1 | valid |
| TWNK | 39780253 | Llama 3.1, Qwen 3, DeepSeek R-1 | valid |

**Table 7.** Manual Validation for Phenotype: Childhood onset sensorineural hearing impairment

| Gene(s) | Supporting PMID | LLM(s) | Validation |
|---|---|---|---|
| BAX | 35815259, 38952359 | Llama 3.1, Qwen 3, DeepSeek R-1 | valid |
| BCL2, CASP3, IL6 | 35815259 | Llama 3.1, Qwen 3 | valid |
| CCDC69, CLMP, FAM110B, PLEKHO1 | 32859214 | Llama 3.1, Qwen 3, DeepSeek R-1 | valid |
| CLDN23, CPT2, CXCL1, ITLN1, LZTS3, TIMP1 | 31893510 | Llama 3.1, Qwen 3 | valid |
| FOXO4 | 32764986 | Llama 3.1, Qwen 3 | valid |
| IL1B | 38952359 | Llama 3.1, Qwen 3, DeepSeek R-1 | valid |
| MAN1B1 | 40264753 | Qwen 3, DeepSeek R-1 | valid |

**Table 8.** Manual Validation for Phenotype: Colon cancer

| Gene(s) | Supporting PMID | LLM(s) | Validation |
|---|---|---|---|
| CCDC34, DNPH1, LILRA5, PURG, RNF207, SEC31B, TRPC6, ZCRB1 | 38097166 | Llama 3.1, Qwen 3 | invalid |
| CD79B, DDHD1, FKBP1B, STXBP4, TRAM2 | 40593136 | Llama 3.1, Qwen 3 | valid |
| LMNA, LMNB1, LMNB2 | 34530169 | Llama 3.1, Qwen 3, DeepSeek R-1 | valid |
| PRAMEF12 | 38913622 | Llama 3.1, Qwen 3, DeepSeek R-1 | valid |

**Table 9.** Manual Validation for Phenotype: Malignant neoplasm of the central nervous system

| Gene(s) | Supporting PMID | LLM(s) | Validation |
|---|---|---|---|
| BCAN, CHMP4C, DOC2A, HPS1, SLC29A3, SLC2A8 | 40751887 | Llama 3.1, Qwen 3, DeepSeek R-1 | valid |
| CD276, CLDN14, ITGA4, L1CAM, NR-CAM, PDCD1, SDC1, SELE, SELPLG | 35117360 | Llama 3.1, Qwen 3, DeepSeek R-1 | valid |
| FOXM1 | 38352275 | Llama 3.1, DeepSeek R-1 | invalid |
| RUNX1 | 40115757 | Llama 3.1, Qwen 3, DeepSeek R-1 | valid |
| VCAM1 | 35117360 | Llama 3.1, Qwen 3 | valid |

**Table 10.** Manual Validation for Phenotype: Myeloid leukemia

| Gene(s) | Supporting PMID | LLM(s) | Validation |
|---|---|---|---|
| AOC3, IDO1, PTGIS | 37582873 | Llama 3.1, Qwen 3, DeepSeek R-1 | valid |
| CD2, COL5A2, ICOS, TAP1 | 32294292 | Llama 3.1, Qwen 3, DeepSeek R-1 | invalid |
| CDK12 | 32172432 | Llama 3.1, Qwen 3, DeepSeek R-1 | invalid |
| CDKN1A, CYP1B1, DKK1, NTS | 37615851 | Llama 3.1, Qwen 3, DeepSeek R-1 | valid |
| ELAVL2 | 38654363 | Llama 3.1, Qwen 3 | valid |
| GDF15 | 38860553 | Llama 3.1, DeepSeek R-1 | invalid |
| L1CAM | 31952346 | Llama 3.1, Qwen 3, DeepSeek R-1 | valid |
| NF1, STK11 | 32733558 | Qwen 3, DeepSeek R-1 | valid |

**Table 11.** Manual Validation for Phenotype: Ovarian neoplasm
